# Complete reconstruction of the unbinding pathway of an anticancer drug by conventional unbiased molecular dynamics simulation

**DOI:** 10.1101/2020.02.23.961474

**Authors:** Farzin Sohraby, Mostafa Javaheri Moghadam, Masoud Aliyar, Hassan Aryapour

## Abstract

Understanding the details of unbinding mechanism of small molecule drugs is an inseparable part of rational drug design. Reconstruction of the unbinding pathway of small molecule drugs, todays, can be achieved through molecular dynamics simulations. Nonetheless, simulating a process in which a drug unbinds from its receptor demands lots of time, mostly up to several milliseconds. This amount of time is neither reasonable nor affordable; therefore, many researchers utilize various biases that there are still many doubts about their trustworthiness. In this work we have utilized short-run simulations, replicas, to make such time-consuming process cost effective. By replicating those snapshots of the trajectories which, after careful analyses, were selected as potential candidates we increased our system’s efficiency considerably. As a matter of fact, we have implemented a sort of human bias, inspecting trajectories visually, to achieve multiple unbinding events. We would like to call this stratagem, replicating of potent snapshots, “rational sampling” as it is, in fact, benefiting from human logic. In our case, an anticancer drug, the dasatinib, completely unbounded from its target protein, c-Src kinase, in only 392.6 ns, and this was gained without applying any internal biases and potentials which can increase error level. Thus, we achieved important structural details that can alter our viewpoint as well as assist drug designers.

## Introduction

Residence time of small molecule drugs in a binding pocket and detailed process of the unbinding pathway are of high importance in the context of rational drug design. Molecular dynamics (MD) simulation has provided a great opportunity to study these processes in details (*1–3*). However, this approach is noticeably time consuming if one wants to map the whole unbinding process. Over the past years, a number of methods have been developed and tested to overcome this obstacle (*4–20*). Most of these studies have made their ways through implementation of internal biases.

Having said that, it is taken for granted that the proteins’ structures are so sensitive to changes, even slight ones, including the pH, temperature, pressure and so on. Therefore, the existence of biases in a simulation inevitably affects ligand’s and protein’s natural motions and consequently their behaviors. It can also affect residues of the unbinding pathway and influence their interactions and causes the ligand to unbind smoothly from the binding site without the necessary conformational changes being made on the protein structure for leaving ligand from one state to another. Hence, if one is supposed to study an unbinding pathway, they should be aware about these undesirable impacts. In this article, we have taken the advantages of the conventional MD to achieve an unbinding event in which an anticancer drug, the dasatinib, unbinds from its protein, c-Src kinase, without using any internal biases or pseudo forces. Thus, our results are more reliable than these other methods. Herein, by simulating ten protein-ligand systems, with duration of 0.5 microseconds per each one and total runtime equal to 5 microseconds, we explored the binding pocket in order to find the key elements which have pivotal roles in the unbinding pathway. Then, we picked up the potential snapshots which possessed our desired criteria to extend them for further short-run simulations i.e. replicas. We will refer to this type of sampling as the “rational sampling” further on.

## Methods

### Molecular dynamics simulation protocol and analyses

All optimizations and binding simulations were initiated without or with pre-equilibrated state of the relevant apo-protein using the OPLS force field (*21*) in GROMACS 2018 (*22*), respectively. For binding simulations, first, the related apo-protein was placed in the center of a triclinic box with a distance of 1.5 nm from all edges, and sixteen relevant ligands were inserted into each simulation box with random positions and solvated with TIP3P water model (*23*). Then, sodium and chloride ions were added to produce a neutral physiological salt concentration of 150 mM and the overall systems had approximately 40000 atoms. Each system was energy minimized, using steepest descent algorithm, until the Fmax was found to be smaller than 10 kJ.mol^−1^.nm^−1^. All of the covalent bonds were constrained using the Linear Constraint Solver (LINCS) algorithm (*24*) to maintain constant bond lengths. The long-range electrostatic interactions were treated using the Particle Mesh Ewald (PME) method (*25*) and the cut off radii for Coulomb and Van der Waals short-range interactions was set to 0.9 nm for Dasatinib-c-Src systems. The modified Berendsen (V-rescale) thermostat (*26*) and Parrinello–Rahman barostat (*27*) respectively were applied for 100 and 300 ps to keep the system in the stable environmental conditions (310 K, 1 Bar). Finally, simulations were carried out under the periodic boundary conditions (PBC), set at XYZ coordinates to ensure that the atoms had stayed inside the simulation box, and the subsequent analyses were then performed using GROMACS utilities, VMD (*28*) and USCF Chimera, and also the plots were created using Daniel’s XL Toolbox (v 7.3.2) add-in (*29*). The free energy landscapes were rendered using Matplotlib (*30*). In addition, to estimate the binding free energy we used the g_mmpbsa package (*31*). All of the computations of this work were performed on an Ubuntu desktop PC with a [dual-socket Intel(R) Xeon(R) CPU E5-2630 v3 + 2 NVIDIA GeForce GTX 1080] configuration. The performance of this platform is ~250 ns/day for running a ~40000-atom system.

## Results and discussion

Entering the binding pocket is a necessity for a drug but not sufficient. After binding, it has to apply the brakes and stays there for a while. In other words, adequate residence time and selectivity are the two highlights of a good binder. In this regard, understanding details of an unbinding process with its key elements is of the high importance for designing effective drugs.

In order to find these key elements, we performed five long replicas with a duration time of 500 ns for each protonated and deprotonated forms of dasatinib (Fig. 1c) in complex with c-Src kinase (Fig. 1a), which cumulated to the total run time of 5 µs (Fig. 5). After careful analysis of the trajectories, frame by frame, we found out that the main reason that made the dasatinib to leave the pocket was its fluctuations. These fluctuations were due to the both protein’s dynamics and reciprocal motions derived from water molecules. The more dasatinib jiggled and wiggled, the more interactions between residues of the binding pocket and dasatinib were weakened; thus, the more likely it was for dasatinib to be unbound.

**Figure 1.**
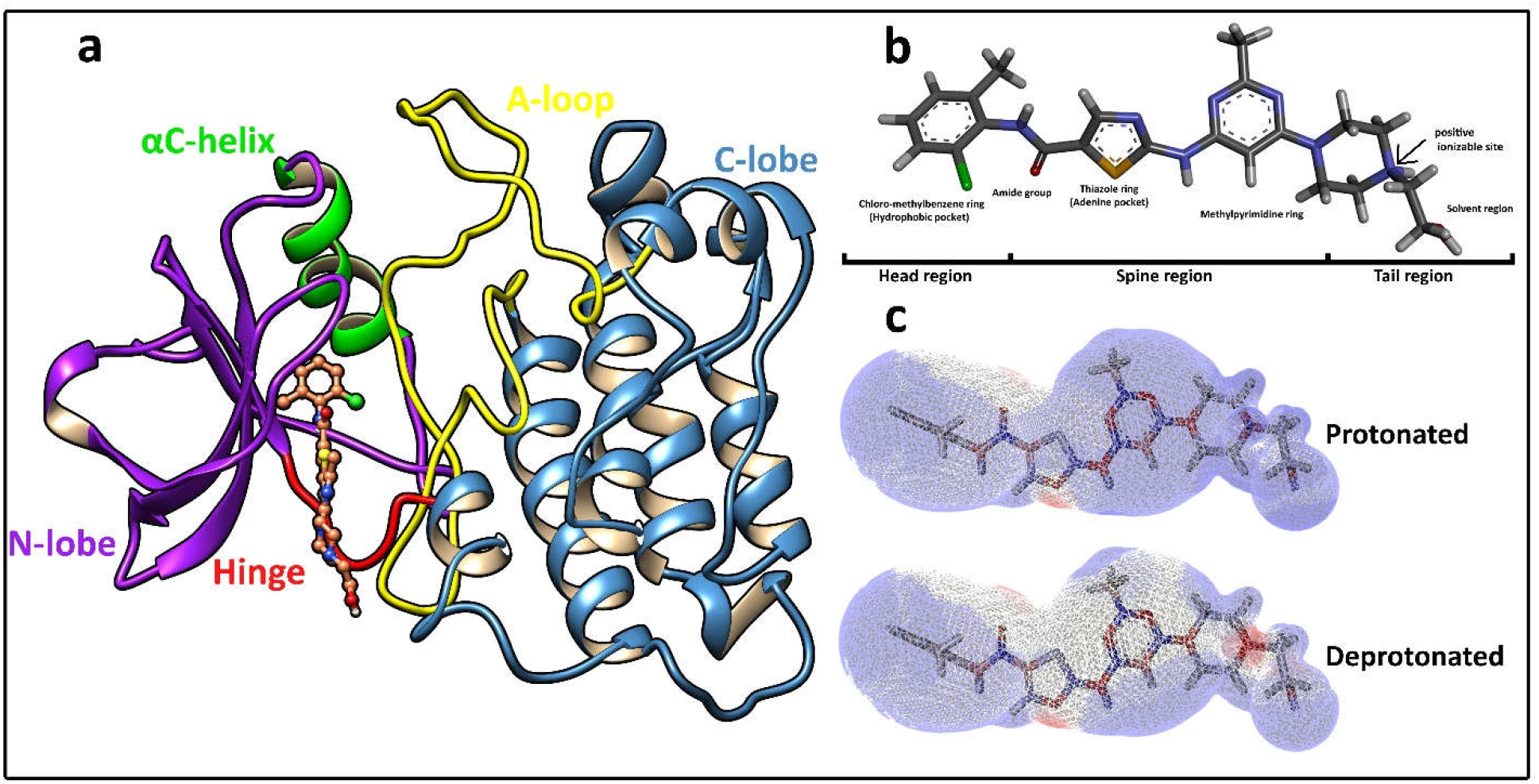
The structures of c-Src and dasatinib. **a**, The crystallographic structure of the c-Src kinase in complex with the dasatinib, and important loops and regions in the structure. **b**, Different functional groups and regions of dasatinib. **c**, The effect of protonation on overall partial charges of dasatinib.

Looking at the c-Src kinase, conformations of two regions have significant impact on the binding pocket. Any changes of the A-loop and the αC-helix can eclipse configuration of the binding pocket. When the A-loop was folded and the αC-helix was at its “in” conformation, bended towards the pocket, more amino acids could interact with the ligand and shield it from water molecules; as a result, this made dasatinib more stable (Fig. 2a). On the other hand, when the A-loop was unfolded and the αC-helix was at its “out” conformation, bended outwards, the binding pocket was fully exposed to water molecules and they were allowed to get inside the binding pocket with ease and so made the ligand unstable. This process brought about more intensive fluctuations (Fig. 2b).

**Figure 2.**
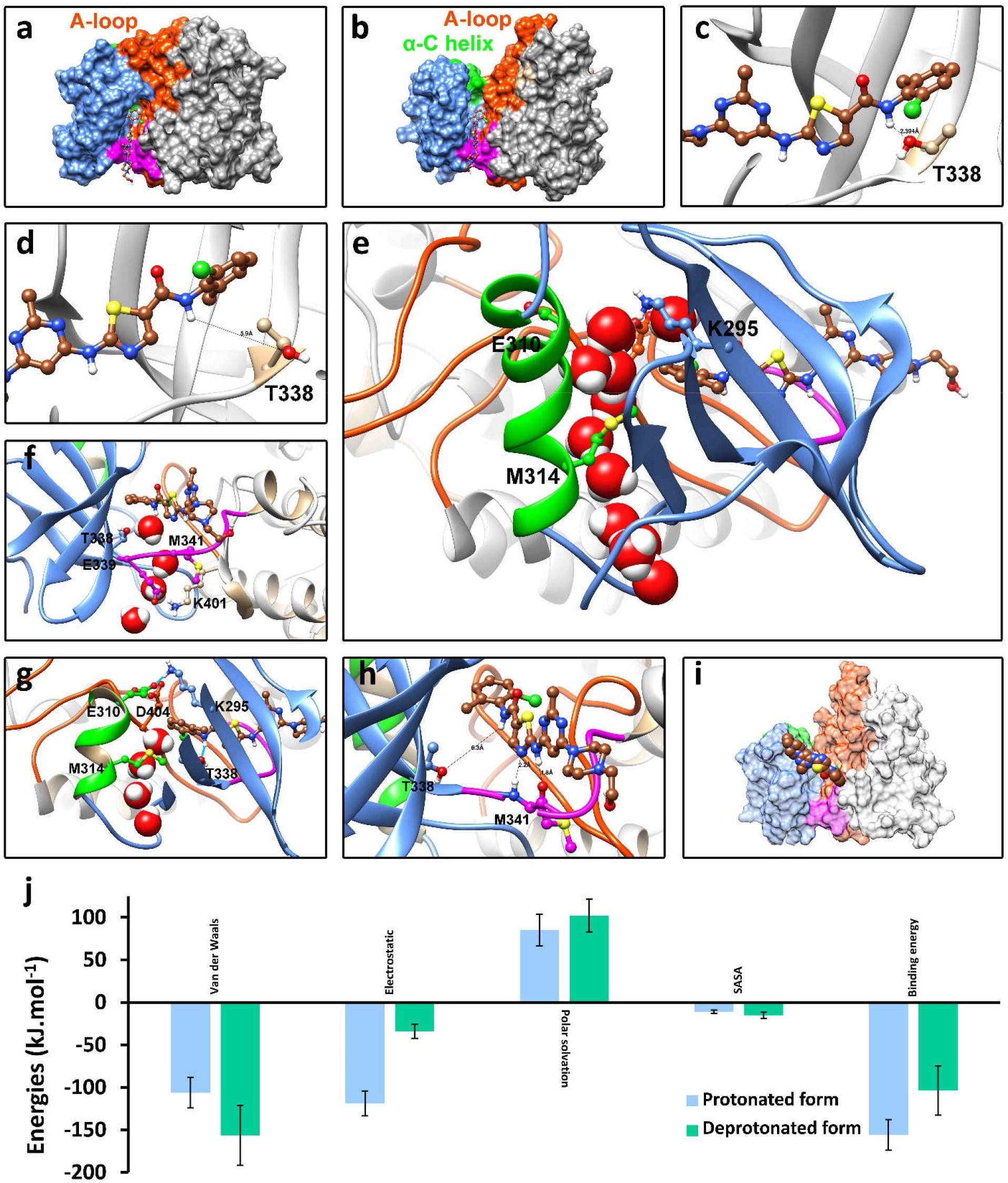
The atomic details of the dasatinib unbinding pathways. **a**, The conformation of c-Src kinase when A-loop was folded and α-C helix was at the “in” conformation. **b**, The conformation of protein when A-loop was unfolded and the α-C helix was at the “out” conformation; the binding pocket was exposed to the solvent molecules. **c**, The established hydrogen bond between OG atom of T338 and dasatinib; it kept the head of dasatinib down, in the “forth state”. **d**, The breakage of hydrogen bond between T338 and dasatinib freed the ligand and it regained the “back state” conformation. This breakage occurred because of dihedral angle rotation of T338 side-chain. **e**, After the salt bridges’ breakage and when the E310 was headed outwards, water channel was created. This channel was mainly under the influence of M314. **f**, Water-mediation of the T338 and dasatinib hydrogen bond was engineered by a single water molecule that was delivered into the pocket by assistance of M341, E339 and K401. These residues were doing somehow like a mouth and transferring (swallowing) water molecules, one by one, inside the pocket. **g**, The role of M314 in delivering water molecules inside the pocket and then conserve them in deep parts of the binding pocket. **h**, The two hydrogen bonds were formed between the backbone atoms of M341 and dasatinib; these bonds kept the spine of dasatinib down and helped the head to be remained at the “back state” conformation. The breakage of these two hydrogen bonds was the rate-limiting step of the unbinding event. **i**, The unbound state of dasatinib. **j**, The average binding free energy of the protonated and the deprotonated dasatinib, in complex with c-Src kinase protein, was calculated during the 5 µs production runs, using the MMPBSA. The values are the mean ± SD.

Periodic breakage of two evolutionary conserved salt bridges, established between K295 and E310/D404, and also interferences of R409 and R419 were the main reasons of the αC-helix-out conformation presence. Nevertheless, although the A-loop plays an important role in both processes of the binding and unbinding, the main cause for its transition, from the folded to the unfolded conformation, is still unknown (*32*).

Some residues that could immensely alter the conformation and orientation of dasatinib inside the binding pocket were detected throughout our simulations. One of these residues, with the most pivotal role, was T338. In the native binding mode, a hydrogen bond, formed between the amide group of the dasatinib’s head segment and the OG atom of the T338 side chain pulled the head of dasatinib towards the T338, presented a state which will be referred as the “forth state” hereinafter (Fig. 2c). Note that different segments and functional groups of the dasatinib are illustrated in figure 1b.

Interestingly, the breakage of this hydrogen bond, under the influence of various factors, made the head of dasatinib to move farther from T338. This state named the “back state” (Fig. 2d). This, in fact, triggered the unbinding event. Generally, there are two main transitions: (i) the transition from the “forth state” to the “back state”, initiated by the breakage of the hydrogen, and (ii) the transition from the “back state” to the complete unbound state, provoked by breakage of two hydrogen bonds which were formed between M341 backbone and the spine of dasatinib.

Recently, Tiwary et al. have studied the nature of the binding pocket of c-Src (*2*). They were of the opinion that as long as the evolutionary conserved salt bridges are in place, water molecules cannot flow inside the binding pocket. Despite their statement, we found that water molecules can flow either inside or outside of the binding pocket with the assistance of some amino acids which from the evolutionary perspective are brilliantly well positioned. Two of them, M314 and M341, were the most important ones. These two residues were able to hand over water molecules to the inner side of binding pocket, with a certain rate. M314 was located at the αC-helix, right under the K295 and E310 salt bridge. Regarding our simulations, M314, most of the time, was in engaged with one or two water molecules through weak hydrogen bonds, even in presence of the salt bridges; it used to hand over water molecules towards the binding pocket, at a low rate (Fig. 2g). However, at some points when the salt bridges were broken and E310 was headed outwards, the M314 channeled water molecules inside the pocket, alongside the αC-helix (Fig. 2e). The consecutive presence of water molecules underneath the head of dasatinib raised the chance of T338 and dasatinib’s hydrogen bond breakage. The M341 was also delivering water molecules inside the pocket, but only M314 was able to channel waters towards the deep parts of the binding pocket. The collaboration of K401, E339 and M341 was seen when a water molecule entered inside the binding pocket, from either side of the K401 and E339 salt bridge. This water molecule used to mediate the T338 and dasatinib hydrogen bond (Fig. 2f). Moreover, rotations of T338 side chain, the dihedral angle exactly, put the OG atom of T338 away from the dasatinib’s amide group and, in row, close to the water molecules (Fig. 2d). All of these incidents led to the breakage of the hydrogen bond which had been formed between T338 OG atom and the nitrogen atom of the amide group. This consequently repositioned the dasatinib’s head segment to the “back state” conformation (Fig. 2h).

There are two hydrogen bonds between the spine of dasatinib and the backbone of M341 which, with the help of surrounding residues, held the spine tightly in the binding pocket (Fig. 2h). In 6 out of 10 long-run simulations the “back state” conformation was observed which means the probability of striking this pose was moderately high (Fig. 5). However, breakage of the two hydrogen bonds, formed between M341 and dasatinib, was considered as a rare event. Having calculated lengths of these two hydrogen bonds, in all of the simulations, only a few snapshots were spotted in which these bonds were either broken or weakened (Fig. 3a, b, c). Surprisingly, none of them lasted more than 2 ns. Thus, it convinced us that these two strong bonds were the main determinates of witnessing a complete unbound state. Thus, the transition from the “back state” to the unbound state can be taken as the rate-limiting step of the unbinding pathway. The free energy landscapes (Fig. 3d, e, f) and the interaction energy shares of each binding pocket residues (Fig. 6) show that the breakage of these two hydrogen bonds, mostly witnessed when the dasatinib were possessing the “back state” conformation, was the rate-limiting step.

**Figure 3.**
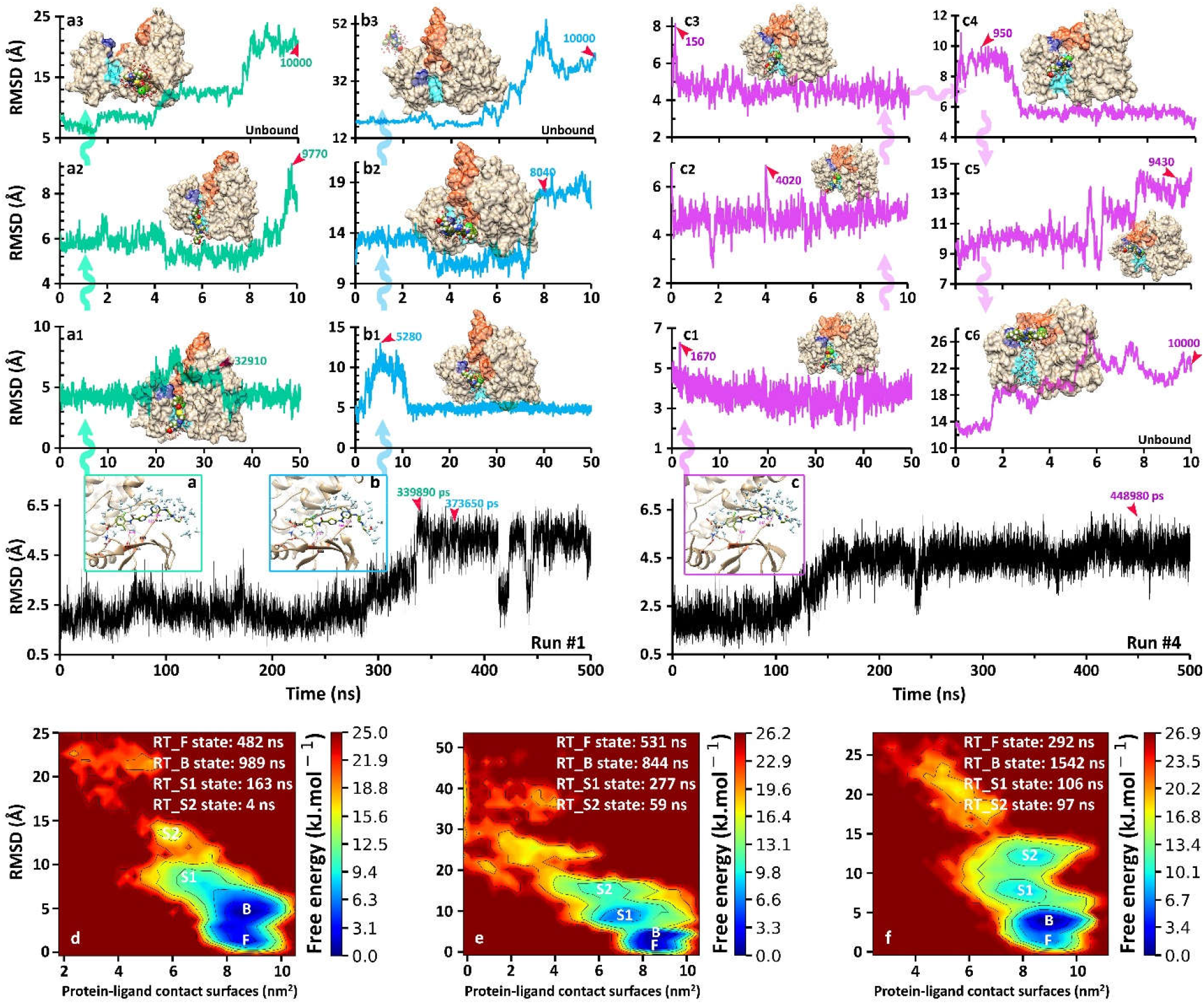
The simulations’ diagrams for unbinding pathways. **a**, The time-frame 339890 ps from the run #1 was picked up. In this frame, dasatinib was at the “back state” conformation and one of the established hydrogen bonds between M341 and dasatinib was weakened. The head of dasatinib was fully solvated. This snapshot was used as the seed for further simulations, consisted of three tandem rounds of simulations (a1 to a3), we named it “A cascade”. After meticulous analysis of each trajectory, the best snapshots were selected. Then each snapshot, indicated by red arrows on the RMSD plots, was replicated for the next round. **b**, The time-frame 373650 from the run #1 was selected. In this frame the dasatinib was at the “back state” conformation and both of the established hydrogen bonds were weakened and the head of dasatinib was fully solvated. The “B cascade” (b1 to b3) was initiated from this snapshot and after achieving a full unbound state it was finished. **c**, The time-frame 448980 ps from the run #4 was pinned. In this frame dasatinib was at the “back state” conformation, and both of the established hydrogen bonds were broken and the spine was water mediated. This snapshot was the original seed for 6 rounds of simulations (c1 to c6), the “C cascade”. This run consumed twice as effort as the run #1, mainly because in this run the A-loop was folded and the α-C helix was located at the “in” conformation, to reach a full unbound state. Note that all of these cascades successful reached full unbound sates. **d**, The free energy landscape of the A cascade, **e**, the B cascade and, **f**, the C cascade. The letters “B”, “F”, “S” and “RT” stand for the “Back state”, “Forth state”, other states and residence time of each state respectively. The hydrogen bonds, blue solid lines, and distances, pink dashed lines, are also presented in a, b and c figures. The surface illustrations represent the binding pocket, G-rich loop and A-loop, with cyan, blue and orange colors, resp.

For the deprotonated dasatinib, two time-frames the run #1, 339890ps (Fig. 3a) and 373650ps (Fig. 3b), and one time-frame of the run #4, 448950ps (Fig. 3c) were selected as the original seeds for next rounds of simulations which were made up of only short-runs (Fig. 4). In each round, good conformations were highlighted and pipelined to the next round, utilizing the “rational sampling”. Eventually, two full unbound states were achieved, with RMSD > 20 Å, from the seeds which were originated from 339890ps and 373650ps snapshots. While three rounds of simulations were sufficient to attain the unbound state for these two, it was more challenging for the other one, the 448950ps snapshot as it demanded 6 rounds of simulations (Fig. 3c1-6). Unlike the run #1, in this run (#4) the A-loop was in its folded conformation. This conformation limits water flows into the pocket and so decreases the intensity of the ligand’s fluctuations. This made the run #4 twice as challenging as the run #1.

**Figure 4.**
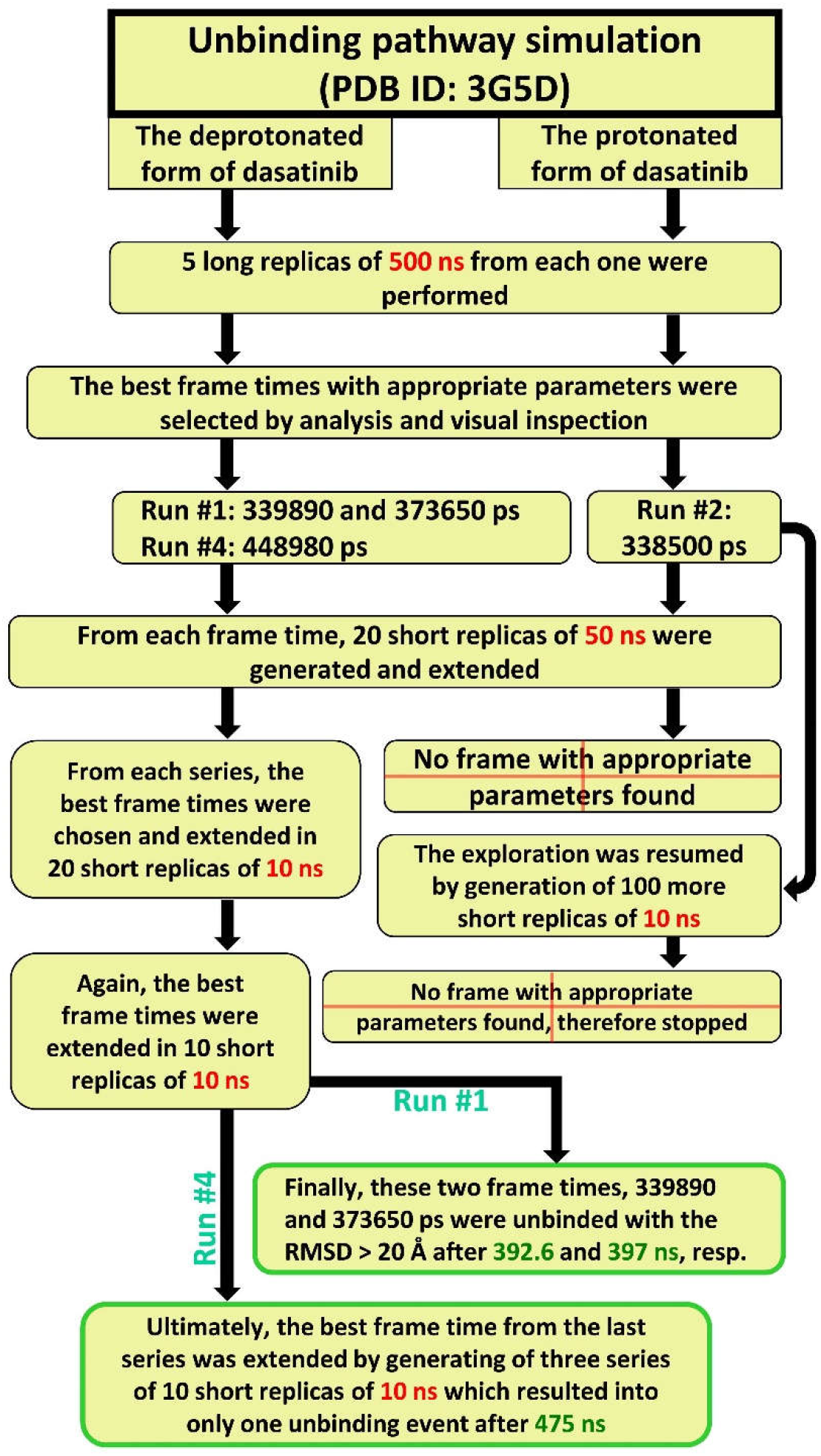
The flowchart of the unbinding pathway.

Our findings also indicate that most of the key events in the unbinding pathway occurred very quick intervals, which is why utilization of short replicas was much more effective for achieving these key events, rather than long-time simulations.

Two different modes were observed through the unbinding. Although all runs possessed similar elements, the sequences of the events were different. In the B and C cascades, once the water molecules seeped underneath the spine, mostly from the tail side, the two hydrogen bonds, established between the spine of dasatinib and M341, were broken, and in its wake the tail and spine were lifted up and solvated. Interestingly, until the last moments of the unbinding, the head was conserving its contacts with the binding pocket.

On the contrary, in the A cascade, it was first the head of dasatinib which was solvated. The water molecules seeped underneath the spine of dasatinib, this time from the inside of the binding pocket, and, after breaking down one of the two hydrogen bonds, they solvated the head segment. The other hydrogen bond was also broken down, but after more simulations. At this point, the ligand was released from the binding pocket and diffused on the protein’s surface.

To estimate the unbinding rate (K_off_) of dasatinib directly, the mean residence time of dasatinib, for these three unbinding events, was equal to 1.9 µs which was far from the experimental observations, 18s (*2*). We opine that this substantial difference was resulted from the rational sampling and the fact that we almost omitted any undesirable simulations. In this respect, the calculation of the K_off_ was a challenge. Although several approaches and formulas have been developed for calculation of the K_off_ by this time (*2, 33, 34*), no one was fitted in our case. Therefore, inevitably the K_off_ should be calculated through statistical method approach which is out of the scope of this article. However, we measured the K_off_ value indirectly using experimentally evaluated dissociation constant (K_d_ = 0.011 µM) (*35*) and our estimated association rate constant (K_on_ = 7.56 s^−1^.µM^−1^), based on previous studies (*36*), which gave the residence time of ~12 s.

Looking at the free energy landscapes of the three unbinding pathways (Fig. 3d, e, f) which were obtained from concatenation of all trajectories of its routes, the stable states were palpable. The graphs represent different unbinding cascades, from a to c. Apart from human biases, neither internal biases nor artificial potentials were implemented in our simulations so the provided figures are more genuine. The analysis of the free energy landscape indicated that the maximum residence time of dasatinib was related to the “Back state”, and the “Forth state” stood at the second place.

Regarding simulations of the deprotonated dasatinib, the “back state” conformation was observable at 22 percent of the total run time, 2.5 µs, while it was 38 percent for protonated form (Fig. 5). Clearly, the protonated form was 1.8 times more inclined than the deprotonated form for being remained at the “back state” conformation. The percentage was calculated by dividing the sum of RMSD values, higher than 3.5 Å, over the total RMSD values. The positive formal charge, existed on the tail of protonated dasatinib, made dasatinib to form much stronger electrostatic interactions with the protein. This, in turn, had a dramatic impact on the entire characteristics of this molecule (Fig. 1c). The binding free energy of the protonated form was lower and its affinity to the pocket was higher, in comparison with the deprotonated form. This, presumably, caused by an elevation in electrostatic interactions (Fig. 2j). Despite the fact that for several times we replicated some potential snapshots of the protonated form, especially when it was at the “back state” conformation, no unbinding was achieved.

**Figure 5.**
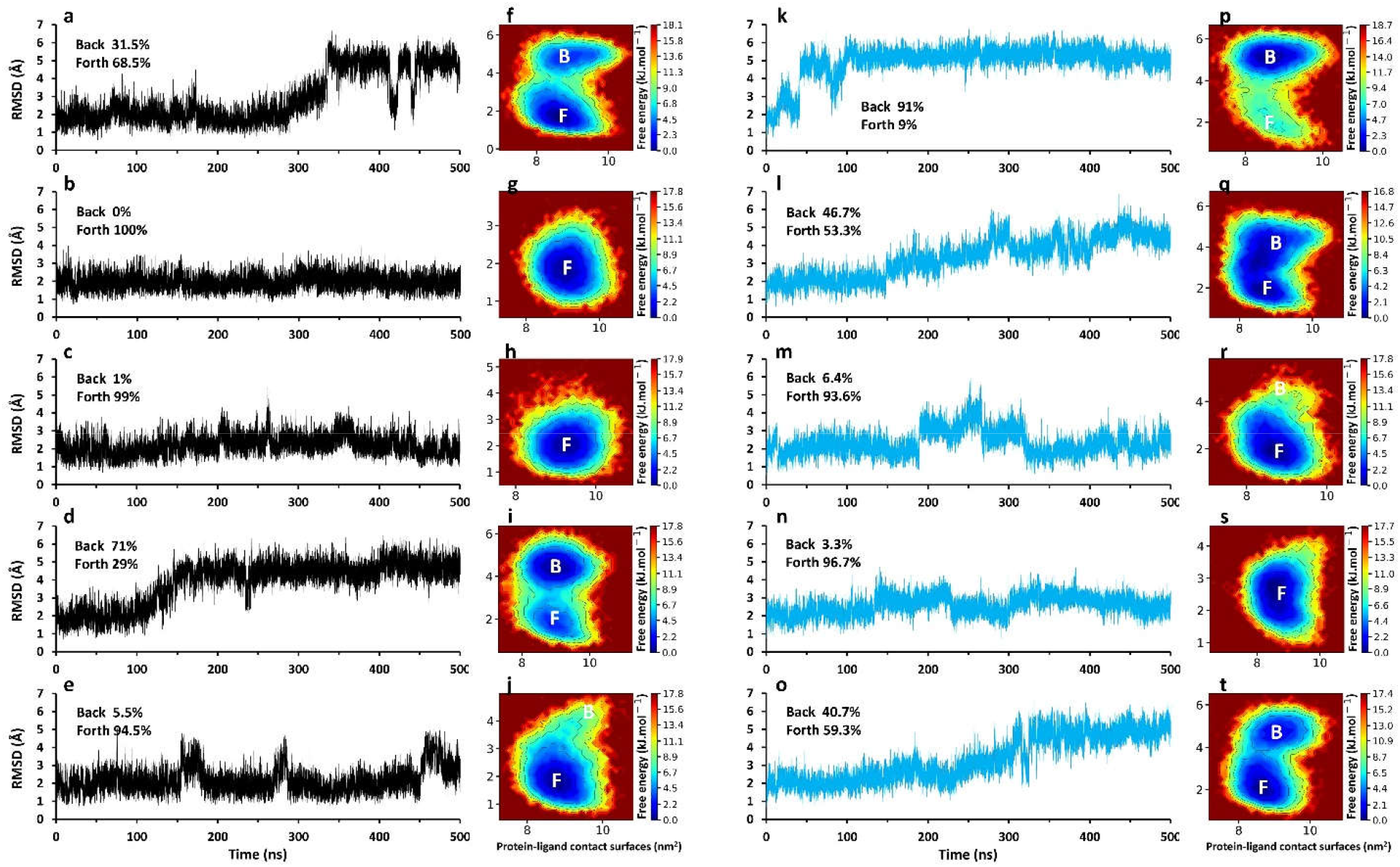
The RMSD plots and the free energy landscapes of the ten first simulations, 500 ns each one. Letter a-j indicate the deprotonated form and k-t depict the protonated form of dasatinib. The letters “B” and “F” stand for the “Back state” and the “Forth state” respectively.

**Figure 6.**
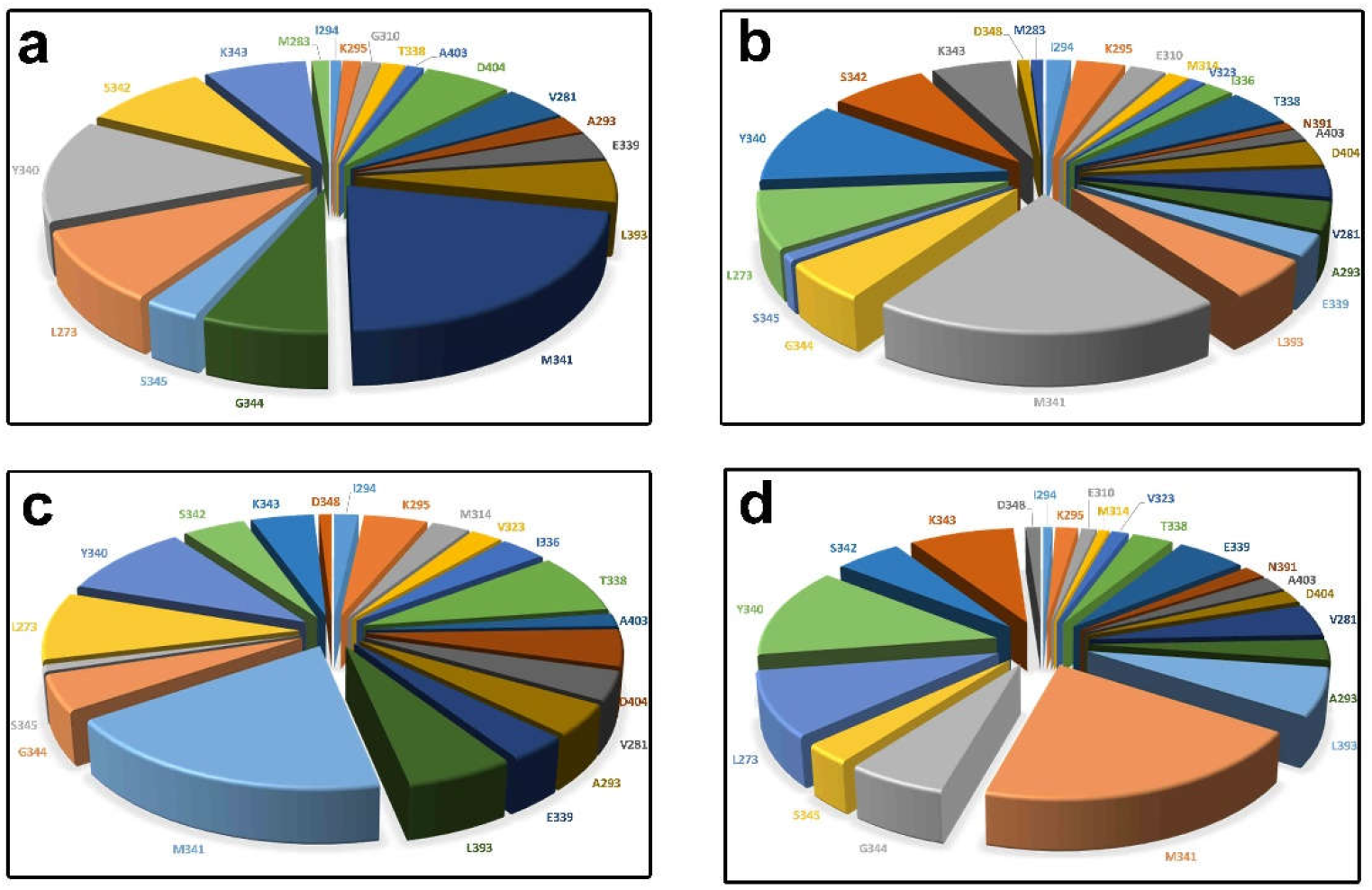
The interaction energy contributions of each residue during the unbinding pathway. **a**, Protonated dasatinib, run#1. **b**, Protonated dasatinib, run#5. **c**, Deprotonated dasatinib, run#2. **d**, Deprotonated dasatinib, run#4.

## Conclusion

Currently, no methods have been able to trace dawn a whole unbinding event of a small molecule drug in real-time. While experimental methods are infeasible in this regard, the MD simulation can be taken workable solution. Because simulating of an unbinding event requires a huge amount of time, micro to milliseconds, the conventional MD simulations i.e. those do not use internal biases, have been considered impractical; therefore, biased methods have been mostly preferred. However, biased methods have their own disadvantages, including false positive results which make these methods rarely trustworthy. In this paper, we have achieved multiple unbinding events with using neither internal biases nor a supercomputer. By utilizing “rational sampling”, constituted short-run simulations as well as human bias, the process of reaching some desirable states and conformations has been speeded up considerably. This is a promising stratagem that empowers researchers to perform such projects with less computational budget. The most paramount point here is that final results would be more reliable than statistical methods and biased-equipped simulations. Thus, drug designers can trust on these results with more comfort. We believe that the details captured in this work can assist drug designers, dealing with kinase inhibitors.

## References

1. P. Tiwary, V. Limongelli, M. Salvalaglio, M. Parrinello, Kinetics of protein–ligand unbinding: Predicting pathways, rates, and rate-limiting steps. Proceedings of the National Academy of Sciences 112, E386 (2015).

2. P. Tiwary, J. Mondal, B. J. Berne, How and when does an anticancer drug leave its binding site? Science Advances 3, (2017).

3. Y. Shan et al., How does a drug molecule find its target binding site? J Am Chem Soc 133, 9181–9183 (2011).

4. M. Perakyla, Ligand unbinding pathways from the vitamin D receptor studied by molecular dynamics simulations. Eur Biophys J 38, 185–198 (2009).

5. Y. Niu, S. Li, D. Pan, H. Liu, X. Yao, Computational study on the unbinding pathways of B-RAF inhibitors and its implication for the difference of residence time: insight from random acceleration and steered molecular dynamics simulations. Phys Chem Chem Phys 18, 5622–5629 (2016).

6. X. Hu et al., Steered molecular dynamics for studying ligand unbinding of ecdysone receptor. Journal of Biomolecular Structure and Dynamics 36, 3819–3828 (2018).

7. Q. Shao, W. Zhu, Exploring the Ligand Binding/Unbinding Pathway by Selectively Enhanced Sampling of Ligand in a Protein–Ligand Complex. The Journal of Physical Chemistry B 123, 7974–7983 (2019).

8. C. Muvva, N. A. Murugan, V. S. Kumar Choutipalli, V. Subramanian, Unraveling the Unbinding Pathways of Products Formed in Catalytic Reactions Involved in SIRT1–3: A Random Acceleration Molecular Dynamics Simulation Study. Journal of Chemical Information and Modeling 59, 4100–4115 (2019).

9. J. Zhu, Y. Lv, X. Han, D. Xu, W. Han, Understanding the differences of the ligand binding/unbinding pathways between phosphorylated and non-phosphorylated ARH1 using molecular dynamics simulations. Scientific Reports 7, 12439 (2017).

10. D. Kosztin, S. Izrailev, K. Schulten, Unbinding of Retinoic Acid from its Receptor Studied by Steered Molecular Dynamics. Biophysical Journal 76, 188–197 (1999).

11. A. Dickson, S. D. Lotz, Multiple Ligand Unbinding Pathways and Ligand-Induced Destabilization Revealed by WExplore. Biophysical Journal 112, 620–629 (2017).

12. J. Rydzewski, O. Valsson, Finding multiple reaction pathways of ligand unbinding. The Journal of Chemical Physics 150, 221101 (2019).

13. A. Nunes-Alves, D. M. Zuckerman, G. M. Arantes, Escape of a Small Molecule from Inside T4 Lysozyme by Multiple Pathways. Biophys J 114, 1058–1066 (2018).

14. P. Carlsson, S. Burendahl, L. Nilsson, Unbinding of Retinoic Acid from the Retinoic Acid Receptor by Random Expulsion Molecular Dynamics. Biophysical Journal 91, 3151–3161 (2006).

15. D. Zhang, J. Gullingsrud, J. A. McCammon, Potentials of Mean Force for Acetylcholine Unbinding from the Alpha7 Nicotinic Acetylcholine Receptor Ligand-Binding Domain. Journal of the American Chemical Society 128, 3019–3026 (2006).

16. A. M. Capelli, G. Costantino, Unbinding Pathways of VEGFR2 Inhibitors Revealed by Steered Molecular Dynamics. Journal of Chemical Information and Modeling 54, 3124–3136 (2014).

17. A. Suresh, A. Hung, Molecular simulation study of the unbinding of α-conotoxin [ϒ4E]GID at the α7 and α4β2 neuronal nicotinic acetylcholine receptors. Journal of Molecular Graphics and Modelling 70, 109–121 (2016).

18. R. O. Dror et al., Pathway and mechanism of drug binding to G-protein-coupled receptors. Proceedings of the National Academy of Sciences 108, 13118 (2011).

19. Y. Niu, D. Pan, Y. Yang, H. Liu, X. Yao, Revealing the molecular mechanism of different residence times of ERK2 inhibitors via binding free energy calculation and unbinding pathway analysis. Chemometrics and Intelligent Laboratory Systems 158, 91–101 (2016).

20. L. Le, E. H. Lee, D. J. Hardy, T. N. Truong, K. Schulten, Molecular Dynamics Simulations Suggest that Electrostatic Funnel Directs Binding of Tamiflu to Influenza N1 Neuraminidases. PLOS Computational Biology 6, e1000939 (2010).

21. G. A. Kaminski, R. A. Friesner, J. Tirado-Rives, W. L. Jorgensen, Evaluation and Reparametrization of the OPLS-AA Force Field for Proteins via Comparison with Accurate Quantum Chemical Calculations on Peptides. The Journal of Physical Chemistry B 105, 6474–6487 (2001).

22. M. J. Abraham et al., GROMACS: High performance molecular simulations through multi-level parallelism from laptops to supercomputers. SoftwareX 1-2, 19–25 (2015).

23. W. L. Jorgensen, J. Chandrasekhar, J. D. Madura, R. W. Impey, M. L. Klein, Comparison of simple potential functions for simulating liquid water. The Journal of chemical physics 79, 926–935 (1983).

24. B. Hess, H. Bekker, H. J. C. Berendsen, J. G. E. M. Fraaije, LINCS: A linear constraint solver for molecular simulations. Journal of Computational Chemistry 18, 1463–1472 (1997).

25. T. Darden, D. York, L. Pedersen, Particle mesh Ewald: An N⋅log(N) method for Ewald sums in large systems. The Journal of chemical physics 98, 10089–10092 (1993).

26. G. Bussi, D. Donadio, M. Parrinello, Canonical sampling through velocity rescaling. The Journal of chemical physics 126, 014101 (2007).

27. M. Parrinello, A. Rahman, Polymorphic transitions in single crystals: A new molecular dynamics method. Journal of applied physics 52, 7182–7190 (1981).

28. W. Humphrey, A. Dalke, K. Schulten, VMD: Visual molecular dynamics. Journal of Molecular Graphics 14, 33–38 (1996).

29. D. Kraus, Consolidated data analysis and presentation using an open-source add-in for the Microsoft Excel® spreadsheet software. Medical Writing 23, 25–28 (2014).

30. J. D. Hunter, Matplotlib: A 2D Graphics Environment. Computing in Science & Engineering 9, 90–95 (2007).

31. R. Kumari, R. Kumar, A. Lynn, g_mmpbsa—A GROMACS Tool for High-Throughput MM-PBSA Calculations. Journal of Chemical Information and Modeling 54, 1951–1962 (2014).

32. D. Shukla, Y. Meng, B. Roux, V. S. Pande, Activation pathway of Src kinase reveals intermediate states as targets for drug design. Nature Communications 5, 3397 (2014).

33. L. Mollica et al., Kinetics of protein-ligand unbinding via smoothed potential molecular dynamics simulations. Sci Rep 5, 11539 (2015).

34. P. Tiwary, V. Limongelli, M. Salvalaglio, M. Parrinello, Kinetics of protein-ligand unbinding: Predicting pathways, rates, and rate-limiting steps. Proc Natl Acad Sci U S A 112, E386–391 (2015).

35. M. Getlik et al., Hybrid compound design to overcome the gatekeeper T338M mutation in cSrc. J Med Chem 52, 3915–3926 (2009).

36. F. Sohraby, H. Aryapour, M. J. Moghadam, M. Aliyar, Ultraefficient Unbiased Molecular Dynamics simulation of protein-ligand interactions: How profound yet affordable can it be? bioRxiv, 650440 (2019).

